# A homeostatic gut-to-brain insulin antagonist restrains neuronally stimulated fat loss

**DOI:** 10.1101/2023.10.20.563330

**Authors:** Chung-Chih Liu, Ayub Khan, Nicholas Seban, Nicole Littlejohn, Supriya Srinivasan

## Abstract

In *C. elegans* mechanisms by which peripheral organs relay internal state information to the nervous system remain unknown, although strong evidence suggests that such signals do exist. Here we report the discovery of a peptide of the ancestral insulin superfamily called INS-7 that functions as an enteroendocrine peptide and is secreted from specialized cells of the intestine. INS-7 secretion increases during fasting, and acts as a bona fide gut-to-brain homeostatic signal that attenuates neuronally induced fat loss during food shortage. INS-7 functions as an antagonist at the canonical DAF-2 receptor in the nervous system, and phylogenetic analysis suggests that INS-7 bears greater resemblance to members of the broad insulin/relaxin superfamily than to conventional mammalian insulin and IGF peptides. The discovery of an endogenous insulin antagonist secreted by specialized intestinal cell with enteroendocrine functions suggests that much remains to be learned about the intestine and its role in directing neuronal functions.

## INTRODUCTION

The central nervous system is known to play a major role in governing systemic lipid homeostasis across species^1–3^. It is also now understood that endocrine hormones signal from organs in the periphery to relay fed-and fasted-state information across the body^4, 5^, including to the nervous system^6, 7^. For example, the intestine has emerged as a preeminent sensory and metabolic organ that relays internal state information via gut hormones to the brain and other important organs in the body^8–11^. It is plausible that many additional endocrine signals relaying interoceptive information to the nervous system remain undiscovered. In the roundworm *C. elegans*, the nervous system plays important roles in regulating fat homeostasis in the intestine^12^, the predominant organ in which lipids are stored and metabolized^13^. Several sensory neurons and the circuits in which they operate are known to regulate the rate and extent of fat utilization. Because these neurons detect and respond to distinct sources of sensory information from the environment, it has become possible to ascribe roles for individual sensory modalities in regulating lipid homeostasis. Previous work from our group has shown that oxygen sensing (in the normoxic range) via the URX and BAG neurons^14^, population density sensing via the ADL neurons^15^ and bacterial food sensing via the serotonergic ADF neurons strongly influence lipid stores^16^. These and other studies^17, 18^ have shown that *C. elegans* calibrates lipid metabolism to the sensory environment in order to optimize behavior, physiology and lifespan.

An interesting feature of the *C. elegans* nervous system is that it does not directly innervate the intestine^16, 19^. To address the question of how sensory information from the nervous system is relayed to the intestine, in previous efforts we had conducted a genetic screen that uncovered a brain-to-gut neuroendocrine peptide called FLP-7, the ancestral ortholog of the mammalian tachykinin family of peptides. The FLP-7 neuropeptide signal is detected by the G protein coupled receptor NPR-22, ortholog of the mammalian Neurokinin 2 Receptor (NK2R), which resides in the intestine where it is necessary and sufficient to trigger fat loss^20^. We developed an assay to measure FLP-7 secretion in living animals and showed that FLP-7 is released by neurosecretory cells called ASI, in proportion to fluctuations in the functions of serotonergic and octopaminergic neurons^20^. Additionally, FLP-7 secretion is also modulated by the oxygen sensing URX neurons and the population density sensing ADL neurons (Extended Data Fig. 1a-d). Thus, the FLP-7/NPR-22 neuroendocrine axis represents the final common pathway by which the sensory nervous system relays information to the intestine for the regulation of lipid metabolism^12^.

Over the course of these studies, we contemplated the possibility that the intestine relayed metabolic information back to the nervous system. Tantalizing evidence from us and others suggested that the *C. elegans* intestine communicates with distinct organs^21–24^, but no clear gut-to-brain molecules had been found. One barrier to identifying such molecules was the lack of readouts of neuronal function and output that are amenable to genetic screens. To test whether the FLP-7 secretion assay could prove to be a metabolically-relevant experimental handle that could circumvent these limitations, we measured the extent to which FLP-7 secretion from ASI neurons (hereafter called FLP-7^ASI^) could be modulated by depleting intestinal fat stores. Inactivation of genes across distinct lipid synthesis pathways previously shown to deplete intestinal fats^22, 25, 26^ (*pod-2*, *acyl CoA Carboxylase; sbp-1, Sterol Regulatory Element Binding Protein; elo-2, Fatty Acid Elongase*) resulted in increased steady-state FLP-7^ASI^ secretion (Extended Data Fig. 1e). Prompted by this observation, we undertook an investigation to identify and define the molecular features underlying gut-to-brain information relay.

## RESULTS

### Discovery of INS-7 as a gut peptide

We reasoned that in the context of lipid metabolism, a molecule that relays internal state information might originate in the intestine since it is the major somatic depot for stored lipids in *C. elegans*. To find such a signal, we conducted an intestine-specific RNAi-mediated screen of the family of genes encoding small peptides, for alterations in FLP-7 secretion from ASI neurons (Fig. 1a). The gene encoding a peptide called *ins-7* emerged as the most potent hit from the screen (Fig. 1b,c); a mutant allele *tm1907* that lacks the bulk of the gene including the F peptide domain, and our newly-generated CRISPR null allele *ssr1532* (Fig. 1d-f) recapitulated the effect of the RNAi: absence of *ins-7* increased FLP-7^ASI^ secretion nearly 2-fold (Fig. 1g,h). INS-7 belongs to the broad insulin/relaxin superfamily of peptides; in *C. elegans* there are 40 members (10 in human) that are thought to be distinguished by their developmental stage-and tissue-specific expression patterns^27, 28^. *C. elegans ins-7* has been reported to be one of the few *ins* genes expressed selectively in adults^28^; a second gene called *ins-19* that has adult-specific expression did not yield a phenotype in the FLP-7^ASI^ secretion assay.

**Fig. 1.**
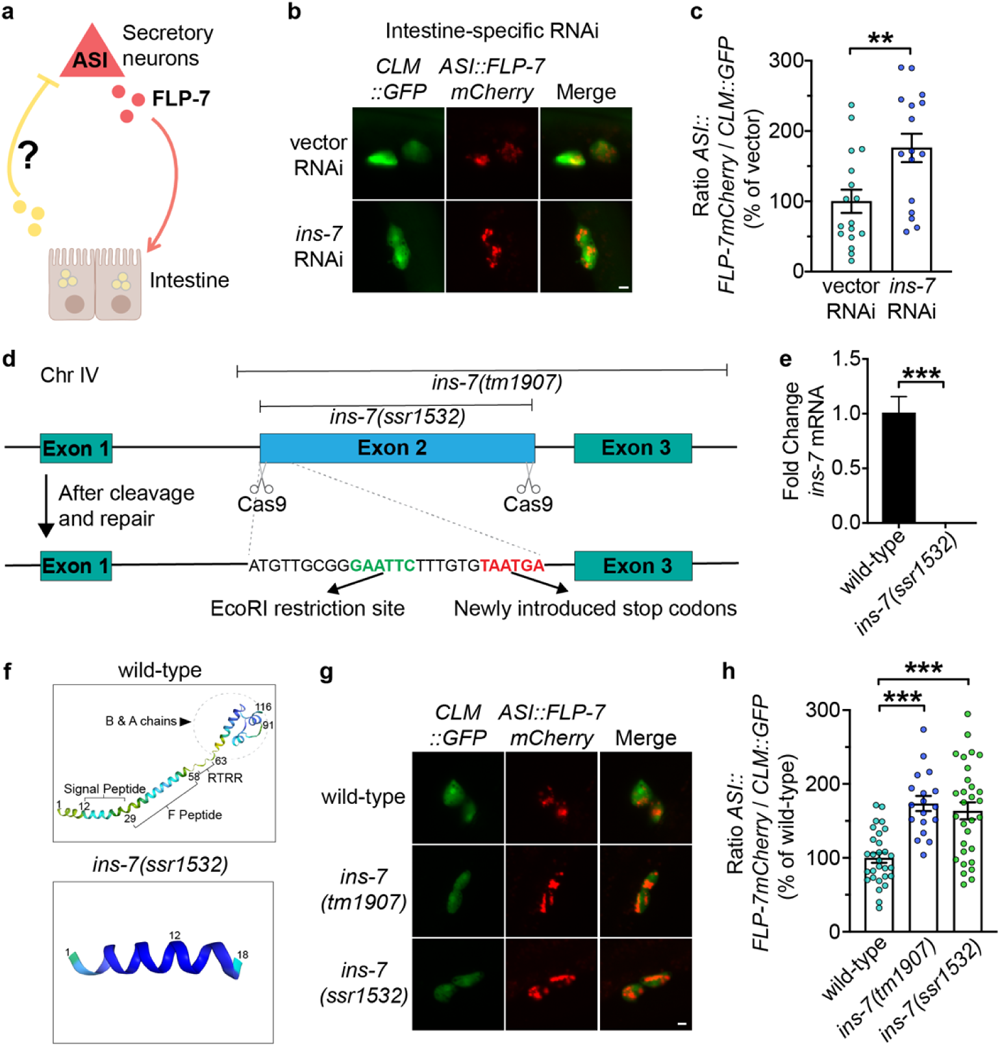
Discovery of *ins-7* as a FLP-7-regulating gene. **a**, Model depicting the *C. elegans* tachykinin neuroendocrine axis and the hypothesized intestine-to-neuron signal. **b**, Representative images of *sid-1; Pvha-6::sid-1* animals bearing integrated *ASI::FLP-7mCherry* and *CLM::GFP* transgenes treated with vector or *ins-7* RNAi. Left panels, GFP expression in coelomocytes; center panels, secreted FLP-7mCherry in coelomocytes; right panels, merge. Scale bar, 5 μm. **c**, The intensity of FLP-7mCherry fluorescence within a single coelomocyte was quantified and normalized to the area of CLM::GFP expression. Assay validation parameters are given in Extended Data Fig 1a-c. Data are expressed as a percentage of the normalized FLP-7mCherry fluorescence intensity of animals treated with vector RNAi ± SEM (n=16-18). **p<0.01 by Student’s *t*-test. **d**, The strategy of CRISPR-Cas9 mediated genome editing, depicting the genomic region of *ins-7* and locations of the Cas9 cut sites for *ins-7(ssr1532)*. Location of introduced EcoRI restriction site (Green) and double stop codons (Red) after cleavage and repair are as indicated. The deletion region of the *ins-7(tm1907)* allele is marked. **e**, qPCR of *ins-7* mRNA in the indicated strains. Pan-actin (*actin-1*, *-3*, *-4*) mRNA was used as a control. Data are presented as fold change relative to wild-type ± SEM (n=3). ***p<0.001 by Student’s *t*-test. **f**, Structure prediction for INS-7 peptides in wild-type or *ins-7(ssr1532)* animals by AlphaFold (DeepMind)^71, 72^. The order of amino acids and specific regions are labeled; the *ins-7(ssr1532)* null allele is a truncation of the *ins-7* gene that results in expression of only the first 18 of 116 amino acids. **g**, Representative images of wild-type, *ins-7(tm1907)* and *ins-7(ssr1532)* animals bearing integrated *ASI::FLP-7mCherry* and *CLM::GFP* transgenes. Left panels, GFP expression in coelomocytes; center panels, secreted FLP-7mCherry in coelomocytes; right panels, merge. Scale bar, 5 μm. **h**, The intensity of FLP-7mCherry fluorescence within a single coelomocyte was quantified and normalized to the area of CLM::GFP expression for each genotype. Data are expressed as a percentage of the normalized FLP-7mCherry fluorescence intensity of wild-type animals ± SEM (n=18-30). ***p<0.001 by one-way ANOVA (Tukey).

An *ins-7::GFP* reporter line we generated revealed robust expression restricted to the first quartet of intestinal cells (Fig. 2a). Called INT1, these anatomically specialized cells define the anterior-most section of the upper intestine, just posterior to the terminal bulb of the pharynx^29^. This intriguingly restricted expression pattern is in stark contrast to the many metabolic genes reported by us and others, which are broadly expressed throughout all the cells of the intestine (INT1-9, 20 cells)^16, 25, 26, 30^. While the intestine-specific RNAi experiment suggested the necessity of *ins-7* expression in INT1 cells, to determine whether *ins-7* expression in INT1 cells is sufficient to rescue the aberrantly-increased FLP-7^ASI^ secretion, we generated transgenic rescue lines in which *ins-7* gene expression was restored in *ins-7(ssr1532)* null mutants under the *ins-7* endogenous promoter, the pan-intestinal promoter *Pvha-6*, as well as a newly-developed INT1-specific promoter, *Pges-1ΔB* (Fig. 2b-e). We found that heterologous *ins-7* expression in INT1 cells alone was sufficient to confer complete rescue of FLP-7^ASI^ secretion to the same extent as restoring *ins-7* expression more broadly across the intestine (Fig. 2b-e).

**Fig. 2.**
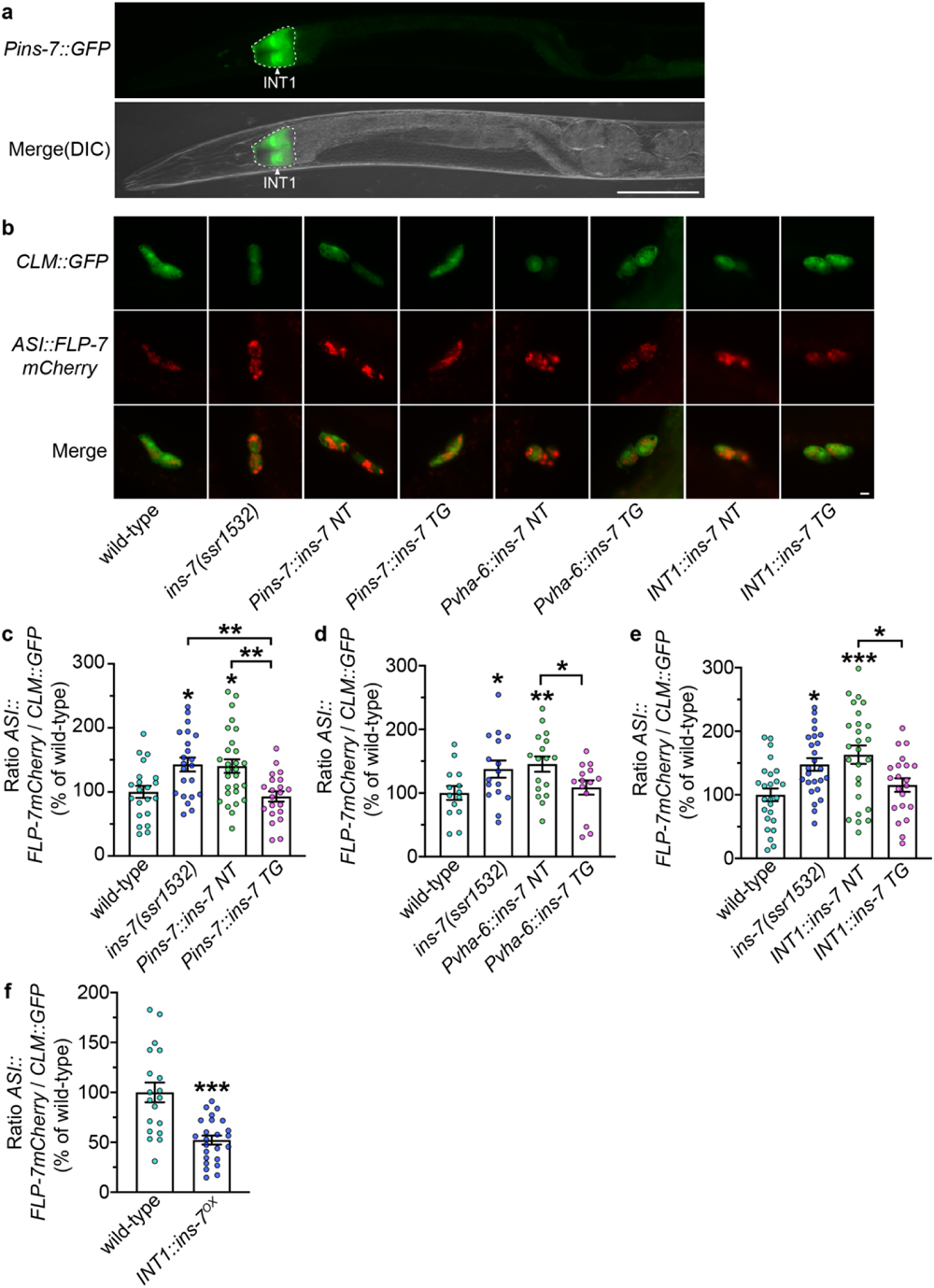
INS-7 functions from the first pair of intestinal cells to regulate FLP-7^ASI^ secretion. **a**, Fluorescent image of a transgenic animal bearing the *Pins-7::GFP* transgene. The white arrowhead indicates GFP expression in INT1 cells. Lower panel, DIC merge; scale bar, 100 μm. **b**, Representative images of wild-type and *ins-7(ssr1532)* animals bearing integrated *ASI::FLP-7mCherry* and *CLM::GFP* transgenes, with the indicated rescuing transgenes. NT, non-transgenic; TG, transgenic. Upper panels, GFP expression in coelomocytes; middle panels, secreted FLP-7mCherry in coelomocytes; lower panels, merge. Scale bar, 5 μm. **c**-**e**, The intensity of FLP-7mCherry fluorescence within a single coelomocyte was quantified and normalized to the area of CLM::GFP expression for each genotype. Data are expressed as a percentage of the normalized FLP-7mCherry fluorescence intensity of wild-type animals ± SEM (n=14-28). *p<0.05, **p<0.01, ***p<0.001 by one-way ANOVA (Sidak). **f**, The intensity of FLP-7mCherry fluorescence within a single coelomocyte was quantified and normalized to the area of CLM::GFP expression for both wild-type and *INT1::ins-7 TG* animals. Data are expressed as a percentage of the normalized FLP-7mCherry fluorescence intensity of wild-type animals ± SEM (n=19-24). ***p<0.001 by Student’s *t*-test.

Further, overexpression of *ins-7* in INT1 cells led to suppression of FLP-7^ASI^ secretion (Fig. 2f), suggesting that INS-7 functions as an instructive cue from the intestine that is necessary and sufficient to drive FLP-7^ASI^ secretion from neurons. INT1 cells were first defined in the context of developmental biology of the *C. elegans* intestine: they have shorter microvilli than the INT2-9 conventional epithelial enterocytes, not unlike the enteroendocrine, goblet and immune M cells of the mammalian intestinal epithelium^29, 31, 32^. Their location as the anterior-most cells of the intestine would also suggest privileged anatomic access to the contents of the gut lumen.

We wished to define additional phenotypes of *ins-7*. In previous studies we had shown that stimulating FLP-7 secretion from ASI neurons activates its cognate receptor NPR-22 in the intestine, whose activity governs fat loss and increases energy expenditure in the intestine via transcriptional activation of the triglyceride lipase ATGL-1^16, 20, 33^. Thus, any condition that increases FLP-7^ASI^ secretion would be predicted to trigger the hydrolysis of fat stored in the intestine via ATGL-1, accompanied by increased energy expenditure. Accordingly, we found that *ins-7* null mutants that display increased FLP-7^ASI^ secretion (Fig. 1g,h), had significantly decreased fat stores (Fig. 3a,b and Extended Data Fig. 2a). This decrease in fat stores was fully dependent on the presence of the *flp-7* gene, because *ins-7;flp-7* mutants suppress the effect of *ins-7* on intestinal fat loss (Fig. 3a,b). The decreased fat stores in the intestine were corroborated by the expected increase in maximal energy expenditure in the *ins-7* mutants as judged by oxygen consumption, which was also dependent on *flp-7* (Fig. 3c,d). As predicted, the decreased intestinal fat stores of *ins-7* nulls were dependent on *atgl-1* induction (Fig. 3e, f), and RNAi-mediated inactivation of *atgl-1* abrogated the *ins-7* fat phenotype (Fig. 3g). The effects of *ins-7* on intestinal fat stores were not dependent on food intake or locomotion, which were not altered in *ins-7* mutants (Extended Data Fig. 2b-d). To corroborate a role for INT1 cells in INS-7-mediated fat loss, we generated an independent set of *ins-7* transgenic rescue lines using the promoters described in Fig. 2 and measured intestinal fat. We observed complete restoration of the decreased intestinal fat stores of *ins-7* null mutants in all cases, again suggesting that *ins-7* functions from INT1 cells to regulate fat stores across the broader intestine (Fig. 3h-j). Finally, overexpression of *ins-7* in INT1 cells resulted in a small but significant increase in fat storage throughout the intestine (Fig. 3k).

**Fig. 3.**
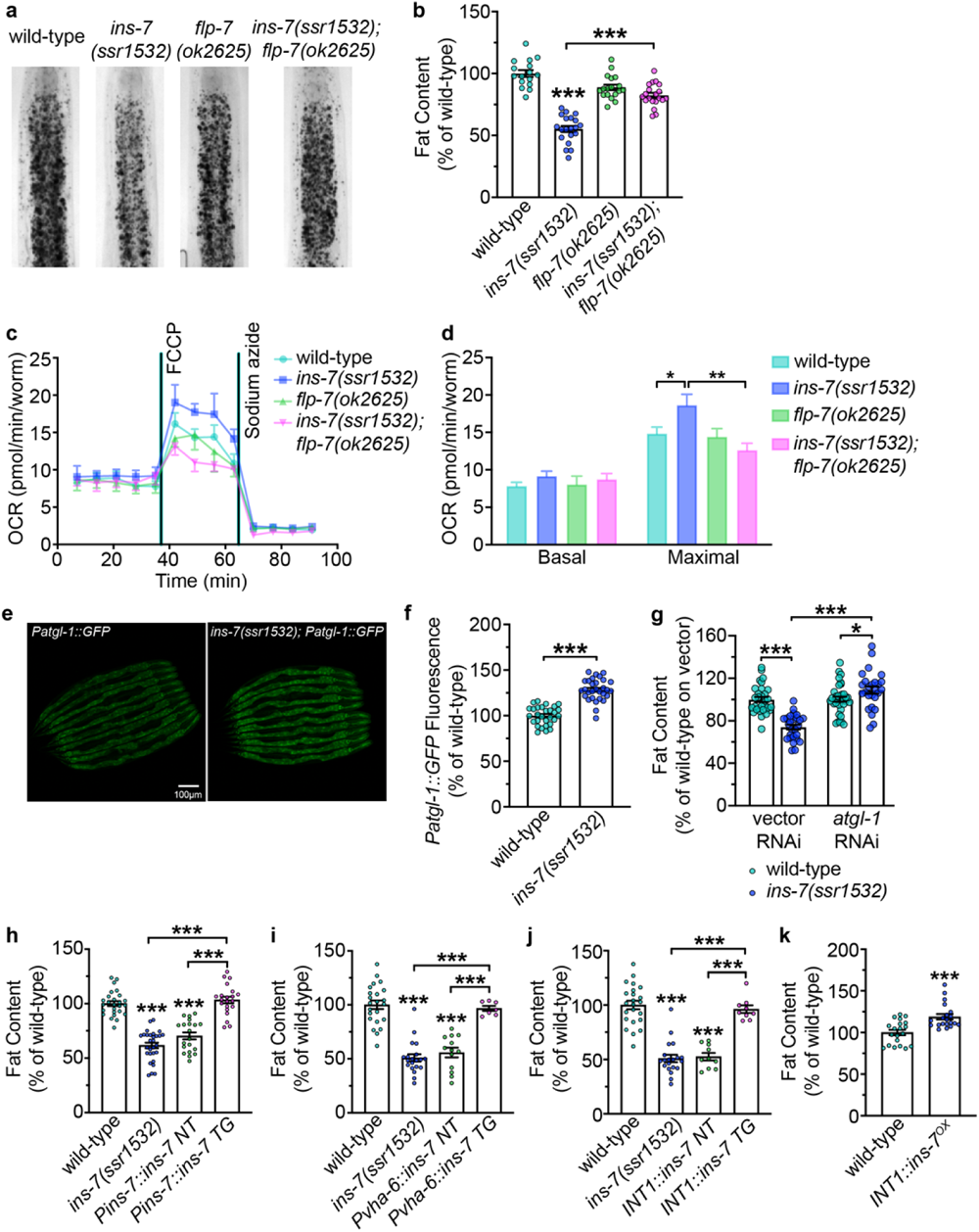
Role of INS-7 in intestinal fat metabolism. **a**, Representative images of wild-type, *ins-7(ssr1532)*, *flp-7(ok2625),* and *ins-7(ssr1532);flp-7(ok2625)* animals fixed and stained with Oil Red O. **b**, Fat content was quantified for each genotype and expressed as a percentage of wild-type animals ± SEM (n=16-20). ***p<0.001 by one-way ANOVA (Sidak). **c**,**d**, Oxygen consumption rate (OCR) of wild-type, *ins-7(ssr1532)*, *flp-7(ok2625)*, and *ins-7(ssr1532);flp-7(ok2625)* animals. Basal OCR was quantified before the addition of FCCP (50mM), and maximal OCR was measured following FCCP stimulation. Data are presented as pmol/min/worm ± SEM (n=16-28 wells, each containing approximately 10 worms). *p<0.05, **p<0.01 by one-way ANOVA (Sidak). **e**, Representative images of wild-type and *ins-7(ssr1532)* animals bearing the integrated *Patgl-1::GFP* transgene. Scale bar, 100μm. **f**, *Patgl-1::GFP* fluorescence was measured and expressed as a percentage of wild-type animals ± SEM (n=29-30). ***p<0.001 by Student’s *t*-test. **g**, Wild-type and *ins-7(ssr1532)* animals treated with vector or *atgl-1* RNAi were fixed and stained with Oil Red O. Fat content was quantified for each genotype and condition and expressed as a percentage of wild-type animals treated with vector RNAi ± SEM (n=26-34). *p<0.05, ***p<0.001 by one-way ANOVA (Sidak). **h**-**j**, Wild-type and *ins-7(ssr1532)* animals with the indicated rescuing transgenes were fixed and stained with Oil Red O. NT, non-transgenic; TG, transgenic. Fat content was quantified for each genotype and expressed as a percentage of wild-type animals ± SEM (n=7-29). ***p<0.001 by one-way ANOVA (Tukey). **k**, Fat content was quantified for both wild-type and *INT1::ins-7^OX^* animals and expressed as a percentage of wild-type animals ± SEM (n=20). ***p<0.001 by Student’s *t*-test.

### Mechanism of INS-7 action

Our data support a model in which *ins-7* functions in the INT1 cells of the intestine to negatively regulate FLP-7^ASI^ secretion from neurons, to regulate fat storage throughout the intestine (Fig. 4a). This model of interorgan communication led us to consider the mechanism of INS-7 action. Phylogenetic analysis of the *ins-7* gene suggests that it belongs to an entirely different clade than that of the conventional mammalian insulin genes, *IGF1* and *IGF2*, which encode proteins that function as agonist ligands for the insulin and IGF receptors^34^ (Fig. 4b). To investigate the mechanism of *ins-7* action, we examined its relationship to *daf-2*, the sole insulin receptor of *C. elegans*^35^. Using the canonical *e1370* allele, we found that *daf-2* mutants had a dramatic reduction in FLP-7^ASI^ secretion, a phenotype in opposition to that of *ins-7*, and *ins-7;daf-2* mutants phenocopied *daf-2* (Fig. 4c,d). Absence of DAF-16/FOXO, the major downstream target of insulin signaling, did not appreciably alter FLP-7^ASI^ secretion, however it completely suppressed the effect of the *daf-2* mutation (Fig. 4c,d). It was possible that the *daf-2* phenotype of reduced FLP-7^ASI^ secretion resulted from *daf-2* expression elsewhere in the nervous system. To address this concern, we conducted antisense inhibition of *daf-2* in ASI neurons alone, which phenocopied the global *daf-2* mutation (Fig. 4e,f), that is, loss of *daf-2*^ASI^ reduced FLP-7^ASI^ secretion. In addition, ASI-specific transgenic rescue of *daf-2* restored FLP-7 secretion to wild-type levels (Fig. 4g,h). Further, intestine-specific RNAi-mediated inhibition of *daf-2* did not alter FLP-7^ASI^ secretion (Fig. 4i,j). Together, these data were most consistent with *ins-7* functioning in opposition to *daf-2*, that is, as an endogenous antagonist to the DAF-2 receptor in ASI neurons. Previous studies of local effects of INS-7 in the nervous system have also defined its action at DAF-2 as antagonistic^36^, although it may function as an agonist in the periphery^37, 38^ (see Discussion).

**Fig. 4.**
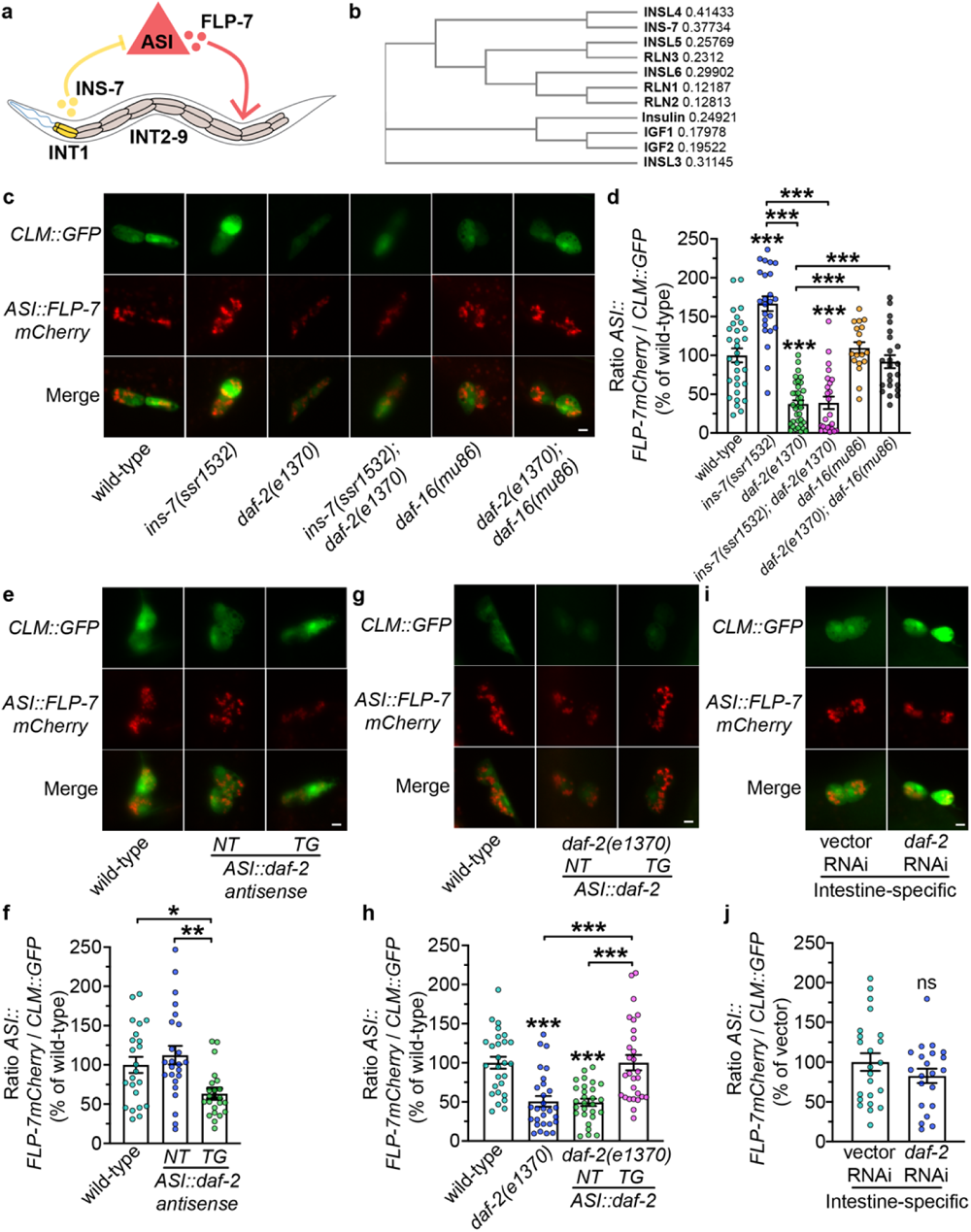
INS-7 functions in opposition to DAF-2. **a**, Model depicting the role of INT1-expressed INS-7 in modulating FLP-7 secretion from ASI neurons. INS-7 is secreted from INT1 cells, which inhibits FLP-7^ASI^ secretion. FLP-7 acts on INT2-9, where the majority of fat is stored. **b**, Phylogenetic tree of INS-7 and members of the human Insulin/Relaxin superfamily by Clustal Omega^73^. The values after each protein name indicate the bootstrap values. **c**, Representative images of wild-type, *ins-7(ssr1532), daf-2(e1370)*, *ins-7(ssr1532);daf-2(e1370)*, *daf-16(mu86)*, and *daf-2(e1370);daf-16(mu86)* animals bearing integrated *ASI::FLP-7mCherry* and *CLM::GFP* transgenes. Upper panels, GFP expression in coelomocytes; middle panels, secreted FLP-7mCherry in coelomocytes; lower panels, merge. Scale bar, 5 μm. **d**, The intensity of FLP-7mCherry fluorescence within a single coelomocyte was quantified and normalized to the area of CLM::GFP expression for each genotype. Data are expressed as a percentage of the normalized FLP-7mCherry fluorescence intensity of wild-type animals ± SEM (n=18-35). ***p<0.001 by one-way ANOVA (Dunnett T3). **e**, Representative images of wild-type FLP-7^ASI^ animals bearing antisense-mediated inactivation of *daf-2* expression in ASI neurons using the *str-3* promoter. NT, non-transgenic; TG, transgenic. Scale bar, 5 μm. **f**, The intensity of FLP-7mCherry fluorescence within a single coelomocyte was quantified and normalized to the area of CLM::GFP expression for each genotype. Data are expressed as a percentage of the normalized FLP-7mCherry fluorescence intensity of wild-type animals ± SEM (n=22-24). *p<0.05, **p<0.01 by one-way ANOVA (Dunnett T3). NT, non-transgenic; TG, transgenic. **g**, Representative images of wild-type and *daf-2(e1370)* animals bearing integrated *ASI::FLP-7mCherry* and *CLM::GFP* with the indicated rescuing transgenes. NT, non-transgenic; TG, transgenic. Scale bar, 5 μm. **h**, The intensity of FLP-7mCherry fluorescence within a single coelomocyte was quantified and normalized to the area of CLM::GFP expression for each genotype. Data are expressed as a percentage of the normalized FLP-7mCherry fluorescence intensity of wild-type animals ± SEM (n=27-28). ***p<0.001 by one-way ANOVA (Dunnett T3). **i**, Representative images of *sid-1; Pvha-6::sid-1* animals bearing integrated *ASI::FLP-7mCherry* and *CLM::GFP* transgenes treated with vector or *daf-2* RNAi. **j**, The intensity of FLP-7mCherry fluorescence within a single coelomocyte was quantified and normalized to the area of CLM::GFP expression for each RNAi treatment. Data are expressed as a percentage of the normalized FLP-7mCherry fluorescence intensity of animals treated with vector RNAi ± SEM (n=21-23). ^ns^p=0.2361 by Student’s *t*-test.

As suggested by our data, we tested the hypothesis that INS-7 acts as an antagonist for neuronal DAF-2 by conducting a direct test of DAF-2 function: we examined DAF-16 localization within the ASI neurons in various genetic contexts. DAF-16 activity as measured by the ratio of its cytoplasmic-to-nuclear localization (abbreviated as C:N ratio) has long been established as an accurate and sensitive hallmark of DAF-2 function^39–41^. In wild-type well-fed animals (in which *ins-7* and *daf-2* are present), DAF-16 resides predominantly in the cytoplasm, such that the DAF-16 C:N ratio is approximately 1.2 (Fig. 5a,b). In *daf-2(e1370)* mutants in which insulin signaling is diminished, as expected, DAF-16/FOXO translocates to the nucleus such that the C:N ratio drops to 0.5 (Fig. 5a,b). However, in the *ins-7(ssr1532)* and *ins-7*^OX^ worms, we also did not observe any deviation in the C:N ratio of DAF-16 localization, relative to wild-type animals (Fig. 5a,b).

**Fig. 5.**
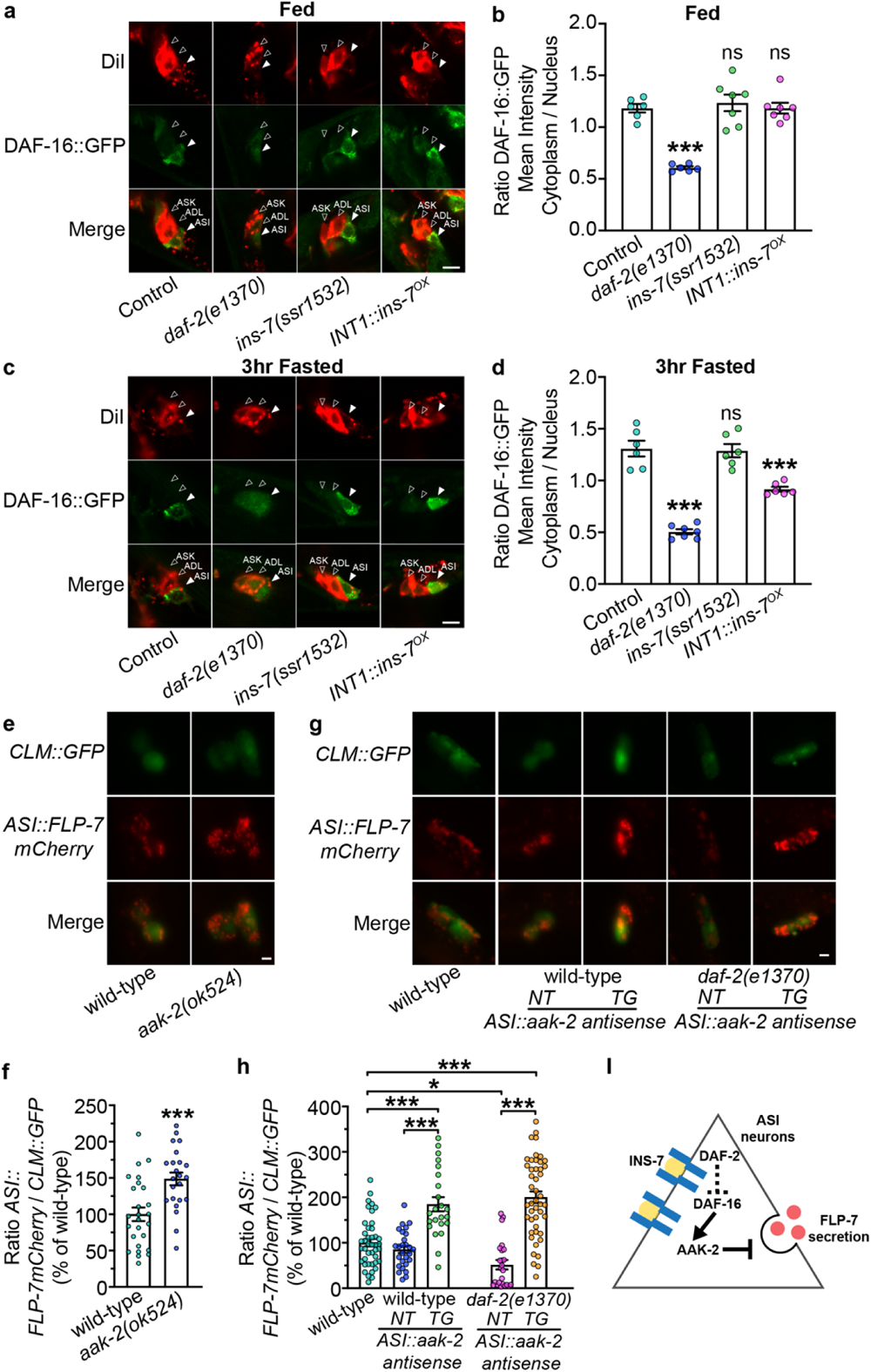
Subcellular DAF-16 localization reveals role for INS-7 as a fasting-induced antagonist. **a**,**c**, Representative images of control, *daf-2(e1370)*, *ins-7(ssr1532)* and *INT1::ins-7^OX^* animals bearing integrated *DAF-16::GFP* transgenes stained with DiI under fed and 3-hour fasted conditions. Upper panels, DiI staining; middle panels, DAF-16::GFP; lower panels, merge. Neurons are indicated with closed arrowheads (ASI) and open arrowheads (ASK and ADL). Scale bar, 5 μm. **b**,**d**, The mean intensity of DAF-16::GFP in the cytoplasm was divided by the mean intensity of DAF-16::GFP in the nucleus for control, *daf-2(e1370)*, *ins-7(ssr1532)*, and *INT1::ins-7^OX^* animals. Data are expressed as a ratio ± SEM (n=6-7). ***p<0.001 by one-way ANOVA (Dunnett). **e**, Representative images of wild-type and *aak-2(ok524)* animals bearing *ASI::FLP-7mCherry* and *CLM::GFP* transgenes. Upper panels, GFP expression in coelomocytes; middle panels, secreted FLP-7mCherry in coelomocytes; lower panels, merge. Scale bar, 5 μm. **f**, The intensity of FLP-7mCherry fluorescence within a single coelomocyte was quantified and normalized to the area of CLM::GFP expression for each genotype. Data are expressed as a percentage of the normalized FLP-7mCherry fluorescence intensity of wild-type animals ± SEM (n=23-26). ***p<0.001 by Student’s *t*-test. **g**, Representative images of wild-type and *daf-2(e1370)* FLP-7^ASI^ animals bearing antisense-mediated inactivation of *aak-2* expression in ASI neurons using the *str-3* promoter. NT, non-transgenic; TG, transgenic. Scale bar, 5 μm. **h**, The intensity of FLP-7mCherry fluorescence within a single coelomocyte was quantified and normalized to the area of CLM::GFP expression for each genotype. Data are expressed as a percentage of the normalized FLP-7mCherry fluorescence intensity of wild-type animals ± SEM (n=23-46). *p<0.05, ***p<0.001 by one-way ANOVA (Sidak). NT, non-transgenic; TG, transgenic. **I**, Model depicting the role of INS-7 as a DAF-2 antagonist in modulating FLP-7 secretion from ASI neurons via DAF-16/FOXO and AMPK signaling.

This result prompted a deeper investigation into the conditions in which *ins-7* normally functions. Conventional insulin, IGF1 and IGF2 are peptides secreted during abundant food availability, and in growth and development. Because the phylogenetic tree suggested that INS-7 belongs to a different clade, and the functional role of INS-7 was in opposition to DAF-2, we postulated that as a physiological antagonist to the insulin receptor, INS-7 may function preferentially in the fasted state, and that its temporal functions may not be fully revealed in well-fed animals. Thus, we tested the effects of a 3-hour fast (a duration sufficient to deplete about 80% of the fat stores in the intestine^22^) on DAF-16 localization. Interestingly, in the fasted condition DAF-16 localization in the ASI neurons did not shift from the previously noted fed state ratios between the cytoplasm and the nucleus in wild-types, *daf-2(e1370)* or *ins-7(ssr1532)* mutants. However, in the fasted state, worms bearing transgenic overexpression of *ins-7* from the INT1 cells (INS-7^OX^) showed a significant shift in DAF-16 localization to the nucleus (C:N ratio 0.8; Fig. 5c,d). This result resembles the effect of the *daf-2(e1370)*-mediated reduction C:N DAF-16 localization (0.5). Thus, the DAF-16^ASI^ localization studies also show that *ins-7* functions in an antagonistic capacity in the context of regulating ASI function and output, and that it functions preferentially in the fasted state.

Multiple lines of evidence show that hundreds of genes containing DAF-16 cis-binding sites function as effectors of the insulin signaling pathway^38, 42–46^. Of these, the AMP-activated protein kinase (AMPK) occupies a central role^47, 48^. As noted in our previous work on the role of ASI neurons in regulating FLP-7 secretion^20^, we found that AMPK α subunit *aak-2(ok524)* mutants had significantly increased FLP-7^ASI^ secretion (Fig. 5e,f). To exclude the possibility that the increased FLP-7^ASI^ secretion in *aak-2* mutants resulted from its absence elsewhere in the nervous system, we used antisense inhibition of *aak-2* solely in the ASI neurons. Antisense inhibition of ASI-specific *aak-*2 resembled the global *aak-2* mutation (Fig. 5g,h), that is, loss of *aak-2*^ASI^ increased FLP-7^ASI^ secretion in a manner indistinguishable from the global absence of *aak-2*. Additionally, ASI-specific antisense inhibition of *aak-2* increased FLP-7^ASI^ secretion in *daf-2(e1370)* mutants to the same extent as it did in the wild-type animals (Fig. 5g,h), thus establishing a mechanistic link between DAF-2, DAF-16 and AMPK in the ASI neurons for the modulation of FLP-7^ASI^ secretion (Fig. 5i). Together, our results show that INS-7 from specialized INT1 intestinal cells inhibits the functional output of ASI neurons to dampen FLP-7 secretion, which drives fat loss throughout the intestine (INT2-9). This effect occurs via INS-7-mediated antagonism of DAF-2 and its downstream actions via DAF-16/FOXO and AMPK signaling in ASI neurons.

### Dynamics of INS-7 function

The experiments delineating a role for INS-7 as a gut-derived signal were designed to reveal steady-state differences in homeostatic mechanisms in well-fed animals. However, our results from the DAF-16 localization experiment suggested that INS-7^INT^^1^ may regulate FLP-7^ASI^ differentially in the fed and fasted states. Additionally, a role for INS-7 as an antagonist might indicate a function in opposition to canonical insulin which is a peptide secreted in the post-prandial state^49^. To test the idea that INS-7 may regulate FLP-7 secretion differentially in the fed and fasted states, we measured the dynamics of FLP-7 secretion in the presence and absence of *ins-7*. In wild-type animals subjected to food deprivation, an increase in FLP-7 secretion is not discernable until 180 minutes (3 hours; Fig. 6a), a time point at which greater than 80% of the intestinal fat stores of the animal have been depleted^22^. Re-exposure to food after 3 hours of fasting restores FLP-7 secretion back to baseline levels within 30 minutes (Fig. 6b), suggesting that the rise in FLP-7 secretion during the fasting regimen is reset by re-feeding. Notably, in *ins-7* null animals (Fig. 6c), the feeding-state-dependent regulation of FLP-7 is abrogated such that FLP-7 secretion levels are chronically high, regardless of feeding or fasting status (Fig. 6c,d) as per the original observation for *ins-7* nulls (Fig. 1). Thus, gut-derived INS-7 provides a brake on FLP-7^ASI^ secretion to inhibit the neuronal drive to trigger fat loss during food shortage.

**Fig. 6.**
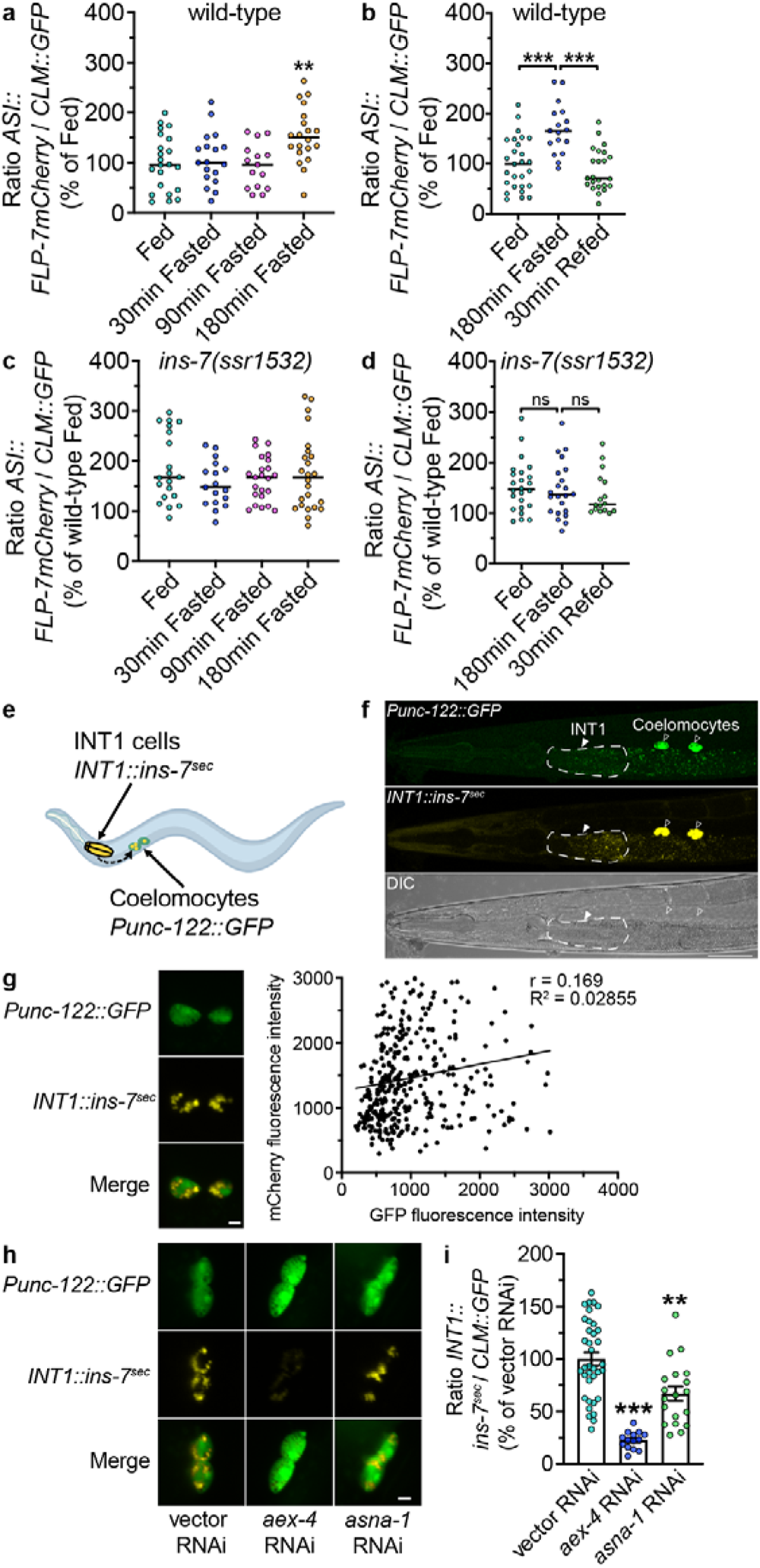
FLP-7^ASI^ secretion dynamics during fasting and re-feeding are controlled by *ins-7*. **a-d**, FLP-7 secretion dynamics during fasting and re-feeding were determined at the indicated time points. For wild-type and *ins-7(ssr1532)* animals, the intensity of FLP-7mCherry fluorescence within a single coelomocyte was quantified and normalized to the area of CLM::GFP expression for each time point. Data are expressed as a percentage of the normalized FLP-7mCherry fluorescence intensity of wild-type fed animals ± SEM (n=15-24). **p<0.01 by by one-way ANOVA (Dunnett). ***p<0.001 by one-way ANOVA (Tukey). **e**, Model depicting the coelomocyte uptake assay for INS-7 secretion. The INS-7mCherry fusion protein (marked in yellow) is expressed in INT1 cells of the intestine, and GFP is expressed in the coelomocytes (marked in green). **f**, Fluorescent image of a transgenic animal bearing *ins-7mCherry* under the control of the INT1-specific promoter. INS-7mCherry signals were observed in INT1 (pseudocolored yellow; closed arrowhead) and coelomocytes (open arrowhead). Upper panel, GFP expression in coelomocytes; middle panel, secreted INS-7mCherry in coelomocytes (pseudocolored in yellow); lower panels, DIC. Scale bar, 50 μm. **g**, mCherry fluorescence intensity values are plotted against GFP fluorescence intensity values for each animal across representative experimental conditions (n=341). Scale bar, 5 μm. **h**, Representative images of *sid-1; INT1::sid-1* animals bearing integrated *INT1::INS-7mCherry* and *CLM::GFP* transgenes treated with vector, *aex-4* or *asna-1* RNAi. Scale bar, 5 μm. **i**, The intensity of INS-7mCherry fluorescence within a single coelomocyte was quantified and normalized to the area of CLM::GFP expression for each RNAi treatment. Data are expressed as a percentage of the normalized INS-7mCherry fluorescence intensity of animals treated with vector RNAi ± SEM (n=14-36). **p<0.01, ***p<0.001 by one-way ANOVA (Dunnett T3).

We investigated whether INS-7 itself is a secreted peptide by generating a transgenic line expressing an INS-7mCherry fusion protein solely from INT1 (pseudo-colored yellow; Fig. 6e,f). To our interest, we found that INT1-expressed INS-7 also accumulates in the coelomocytes, the characteristic hallmark of a secreted peptide in *C. elegans*^20, 50, 51^. Importantly, INS-7 accumulation in coelomocytes does not correlate with their GFP intensity suggesting that these two parameters are independent of one another (Fig. 6g). Extensive validation of the coelomocyte secretion assay by us and others^20, 50^ has shown that it faithfully represents the steady-state levels of a given secreted peptide under a broad range of genetic and physiological conditions. Intestine-specific RNAi of *aex-4*, an intestinal SNARE protein required for vesicular protein secretion^52^ greatly diminished INS-7 secretion, as did *asna-1*, an ATP hydrolase known to specifically regulate insulin secretion^53^ (Fig. 6h,i). Thus INS-7 is a bona fide gut peptide secreted by the INT1 cells of the intestine and acts as a homeostatic signal to restrict FLP-7^ASI^ secretion during food deprivation. Together, our results define a novel gut-brain pathway that relays intestinal state information during acute food shortage.

## DISCUSSION

Here, we report that a peptide of the insulin family called INS-7 is released from specialized INT1 enteroendocrine cells of the *C. elegans* intestine. INS-7 functions as an antagonist at the canonical DAF-2 receptor in the ASI neurons to inhibit FLP-7 secretion. Because FLP-7^ASI^ secretion is a signal that integrates neuronal information to promote fat loss, our results show that the INS-7 gut-to-brain inhibitory peptide serves to limit this signal in the absence of incoming food to the intestine. Work presented here describes the molecular features of our previously postulated hypothesis that homeostatic mechanisms must exist to ensure that neural signals that stimulate fat loss in the intestine are only deployed when there are sufficient fat reserves to do so^22^. These findings uncover a hitherto unknown mechanism of gut-to-brain homeostatic communication in which *C. elegans* lipid metabolism balances between external sensory cues and internal metabolic states.

The identification of INT1 cells as specialized secretory cells of the intestine is intriguing. As the foremost cells of the intestine immediately adjoining the pharynx in the alimentary canal^29^, they have privileged access to incoming nutrient information before food absorption has begun. Sulston *et al* showed that INT1 cells have shorter microvilli than the rest of the intestinal cells (INT2-9)^29^, suggesting that INT1 cells may possess alternate functions in the context of INS-7 secretion that remain to be discovered. Based on our results, we propose that the INT1 quartet functions as enteroendocrine cells that receive information not only from the lumen, but also from the rest of the intestine.

Amongst the many peptides we screened for gut-to-brain signals, INS-7 emerged as the primary suppressor of FLP-7^ASI^ secretion. Initial reports describing a role for INS-7 in longevity regulation had suggested an agonist function at the DAF-2 receptor^37, 38^, whereas later reports suggested antagonistic functions in olfactory learning^36, 54^. It is possible that an explanation lies within the expression features of the various DAF-2 isoforms. In the *C. elegans* nervous system, mutually exclusive expression of DAF-2a and DAF-2c isoforms has been noted. Multiple lines of evidence show that neuronal DAF-2c expression in sensory neurons including the ASI neurons is further induced by fasting and drives the antagonistic effects of DAF-2 signaling in the nervous system^55–57^. Our results indicate a clear role for INS-7 as an antagonist for DAF-2 in ASI neurons; we postulate that agonism and antagonism of insulin peptides at the DAF-2 receptor is determined in a cell-type-specific manner. In contrast to gut-derived INS-7, neuronally-derived INS-7 has been implicated in innate immunity and pathogen avoidance^58^.

Phylogenetic analysis shows that INS-7 may be more orthologous to the mammalian insulin-like peptides and to the relaxin superfamily, rather than the canonical insulin/IGF peptides. Most of the mammalian insulin superfamily members remain poorly characterized, however the insulin-like peptide INSL5 is secreted by L cells of the colon and is a gut hormone thought to be secreted in response to dietary fat intake, and modulates food intake via the hypothalamus^59–61^. The many uncharacterized insulin family peptides across species suggest that new features of interorgan communication regulated by insulin signaling remain to be discovered in additional species. For example, an insulin antagonist may be secreted and function primarily in the fasted state, in opposition to canonical insulin which signals the fed state. We speculate that such a ‘food absence’ signal from the intestine may help animals optimize foraging and food-seeking behaviors while retaining energy reserves during short-term fasts. Finally, the discovery of a specialized intestinal cell with enteroendocrine functions suggests that much remains to be learned about the *C. elegans* intestine and its role in directing neuronal functions.

## METHODS

### *C. elegans* maintenance and strains

Worms were cultured on nematode growth medium (NGM) agar plates with *Escherichia coli* strain OP50 at 20°C as described^62^. The N2 Bristol strain was obtained from the *Caenorhabditis* Genetic Center (CGC) and used as wild-type. All mutant and transgenic strains used in this study are listed in Supplementary Table 1. Worms were synchronized for Oil Red O staining and oxygen consumption by hypochlorite treatment, after which hatched L1 larvae were seeded on plates with the appropriate bacteria; worms were synchronized for neuropeptide secretion assays and behavioral assays by letting gravid adult worms lay eggs on plates with the appropriate bacteria for 1.5 hours. For fasting experiments, young gravid adult worms were transferred with platinum wire without bacteria, to intermediate unseeded NGM plates first after which they were transferred to new unseeded NGM plates for the fasted conditions. All experiments were performed on day 1 adults.

### Cloning and transgenic strain construction

Promoters and cDNAs were amplified using standard PCR techniques from N2 genomic DNA or cDNA and subcloned into expression vectors using Gateway Cloning Technology (Life Technologies). Primers used for cloning in this study are listed in Supplementary Table 2. All transgenic rescue constructs were generated using polycistronic GFP. Transgenic strains were generated by injecting plasmids into the germline of wild-type or mutant worms followed by visual selection of co-injection markers (*Punc-122::GFP* or *Plin-44::GFP*) under the fluorescence microscope. To inhibit the expression of *daf-2* or *aak-2* specifically in ASI neurons, we generated plasmids for antisense mediated inhibition^63^ with the *str-3* promoter. Sense and antisense sequences targeting *daf-2* or *aak-2* were amplified from N2 lysates and subcloned into donor vectors using Gateway Cloning Technology (Life Technologies). The final plasmids for the sense and antisense expression of *daf-2* or *aak-2* under the *str-3* promoter were generated using Gateway Cloning Technology (Life Technologies) and injected into FLP-7 secretion line (SSR1164) at 5 ng/μL (*daf-2*) or 1 ng/μL (*aak-2*) each. The FLP-7 secretion line was previously developed and validated^20^.

For the INS-7 secretion line, N2 worms were injected with 10 ng/μL of the *INT1::ins-7mCherry* plasmid, 15 ng/μL of a *Punc-122::GFP* plasmid, and 75 ng/μL of an empty vector to bring the final concentration of injection mix to 100 ng/μL. A transgenic line with high transmission rate and consistent expression was integrated using the Stratalinker UV Crosslinker 2400 (Stratagene) and backcrossed six times before experimentation. For other microinjections, we injected the animals with 5-25 ng/μL of the desired plasmid, 25 ng/μL of *Punc-122::GFP* or 10 ng/μL of *Plin-44::GFP* and empty vector to maintain a final injection mix concentration of 100 ng/μL. Two lines were selected for experimentation based on the transmission rate and consistency of expression.

### RNAi and qPCR

RNAi experiments were performed as described^64, 65^. Carbenicillin-IPTG plates were seeded with HT115 bacteria containing the empty vector or the relevant RNAi clone and allowed to grow for four days before seeding larvae. Total RNA was extracted from Day 1 adults using TRIzol reagent (Invitrogen). Genomic DNA was isolated using an RNase-free DNase kit (QIAGEN). cDNA was prepared using iScript Reverse Transcription Supermix for RT-qPCR kit (Bio-Rad) according to the manufacturer’s instructions. Quantitative RT-PCR was performed using the SsoAdvanced Universal SYBR Green Supermix (Bio-Rad) following the manufacturer’s instructions. Data were normalized to actin mRNA. Primer sequences used in this study are listed in Supplementary Table 2.

### CRISPR-Cas9 gene editing

Guide RNAs for generating the *ins-7* null allele were designed using the CRISPR guide RNA selection tool^66^. The *dpy-10* guide RNA and repair template were used as reported^67^. All sequences of guide RNAs and repair templates used in this study are listed in Supplementary Table 2. Steps for CRISPR-Cas9 gene editing are briefly described as follows: 0.88 μL tracrRNA (200μM, IDT), 0.82 μL *ins-7* guide RNAs (100μM, IDT), and 0.12 μL *dpy-10* guide RNA (100μM, IDT) were mixed and incubated at 95°C for 5 minutes. 2.52 μL Cas9 protein (IDT, Catalog# 1081058) was added to the mix and incubated at 25°C for 5 minutes. 0.6 μL *ins-7* repair template (100μM, IDT), 0.5 μL *dpy-10* repair template (10μM, IDT), and 3.74 μL nuclease-free water were added to the mix and incubated at 25°C for 60 minutes. The final injection mix was loaded using a pulled capillary needle (1B100F-4, World Precision Instruments) and injected into the germline of N2 young adults. Screening strategy for isolating *ins-7* null mutants was followed as previously described for *C. elegans* CRISPR allele isolation^68^. The final *ins-7* null mutant (*ins-7*(*ssr1532*)) used for experiments was backcrossed four times before experimentation.

### Oil Red O staining

Oil Red O staining was performed as described^16^. Briefly, worms were harvested with phosphate-buffered saline (PBS) and incubated on ice for 10 minutes before fixation. Worms were then stained in filtered Oil Red O (Thermo Scientific) working solution (60% Oil Red O in isopropanol: 40% water) overnight. Approximately 2000 worms were fixed and stained for all genotypes within a single experiment. For each experimental condition, we visually observed about 100 worms on slides, after which 15-20 worms were randomly chosen for imaging. All experiments were repeated at least three times.

### Lipid extraction and quantification

Lipid extraction was performed as described^14^. For each group, 2000 worms per 10cm plate were grown at 25°C until the worms reached young adult stage. After washing with PBS twice, worms were flash frozen in liquid nitrogen. Worms were homogenized in PBS containing 5% TritonX-100 and a protease inhibitor cocktail (Roche), and lipid was extracted using the TissueLyser II (QIAGEN) or a Dounce homogenizer. Triglyceride content was measured using the EnzyChrom Triglyceride Assay Kit (BioAssay Systems), and triglyceride levels were normalized to total protein level determined by the Pierce BCA Protein Assay (Thermo Scientific).

### Oxygen consumption

Oxygen consumption rates (OCR) were measured using the Seahorse XFe96 Analyzer (Agilent) as described^14^. Adult worms were washed with M9 buffer and transferred into a 96-well plate with approximately 10 worms per well. Five basal measurements were taken and then FCCP (50 mM) was injected into each well to measure maximal OCR. Lastly, sodium azide (40 mM) was injected to measure residual OCR. Values were normalized to the number of worms per well. Basal OCR was the average of all measured values prior to the addition of FCCP (50 mM); maximal OCR was the average of the first two measured values after FCCP injection.

### Food intake

Food intake was measured by counting pharyngeal pumping, as described^65^. For each worm, the rhythmic contractions of the pharyngeal bulb were counted over 10 s under a Zeiss M2 Bio Discovery microscope. For each genotype, 25 worms were assessed, and the experiment was repeated at least three times.

### Enhanced slowing response

The enhanced slowing response was measured as described^69^. Day 1 adults were washed off OP50 plates with PBS, washed five times to remove bacteria and placed on NGM agar plates without OP50. After 30 min off food, worms were collected with PBS and seeded onto NGM plates with OP50. Worms were allowed to acclimatize for 5 min, after which the number of body bends per 20 s was counted.

### Thrashing assay

Thrashing rate was measured as described^70^. For each worm, a movement in which the head and/or tail swung to the other side and back to the original position was counted as one thrash. 15-20 Day 1 adults were assessed for each phenotype.

### Image acquisition and quantitation

Oil Red O-stained worms were imaged using 10X objective on a Zeiss Axio Imager microscope. Images were acquired with the Software AxioVision (Zeiss). Lipid droplet staining over the whole body of each worm was quantified using ImageJ software (NIH). All reported results were consistent across biological replicates. Fluorescent images of reporters for FLP-7 and INS-7 secretion were acquired with software AxioVision (Zeiss) using a 20X objective on a Zeiss Axio Imager microscope. The first pair of coelomocytes was imaged. mCherry fluorescence intensity in one of the two imaged coelomocytes was quantified and normalized to the surface area of the coelomocyte (*unc-122::GFP*) as previously described and validated^20^. Within each experiment, at least 15 worms from each condition were quantified using ImageJ software (NIH). Fluorescent images of *ins-7* expression pattern (*Pins-7::GFP*) were collected with software AxioVision (Zeiss) using a 10X objective on a Zeiss Axio Imager microscope. Fluorescent images of DAF-16::GFP and DiI-stained neurons were collected with software NIS-Elements (Nikon) using a 60X objective on a Nikon A1 confocal microscope.

### Statistics

Wild-type animals were included as controls for every experiment. Error bars represent the standard error of the mean (SEM). Student’s t-test, one-way ANOVA, and two-way ANOVA were used as indicated in the figure legends. All statistical analyses were performed using GraphPad Prism 9.3 (GraphPad Software). Appropriate multiple comparison corrections were used for ANOVAs.

## ACKNOWLEDGEMENTS

This work was supported by research grants to S.S. from the NIH/NIDDK (R01 DK124706) and NIH/NIA (R01 AG056648). Strains were provided by the National Bioresource Project (Japan) and the *Caenorhabditis* Genetics Center, which is funded by NIH Office of Research Infrastructure Programs (P40 OD010440). We thank Dr. Anthony Perez, Ms. Aayushi Shah and Ms. Cassandra White of the Srinivasan lab for their critical comments on the manuscript. C.L. was supported by a Dorris Scholar Award from the Dorris Neuroscience Center, The Scripps Research Institute. Elements of Figs. 1a, 6e and Extended Data Fig. 1a were created with BioRender.

---

**Extended Data Fig. 1.**
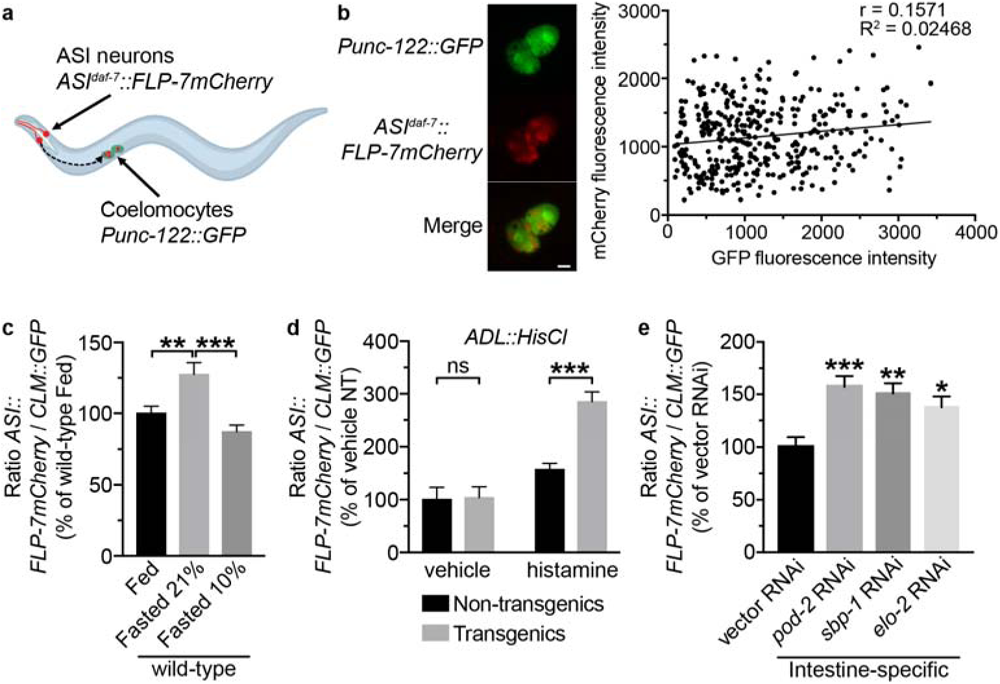
**a**, Model depicting the coelomocyte uptake assay for FLP-7 secretion. The FLP-7mCherry fusion protein (marked in red) is expressed in ASI neurons, and GFP is expressed in the coelomocytes (marked in green). The ratio of red:green fluorescence is used to quantify the level of FLP-7 secretion under different experimental conditions. **b**, FLP-7mCherry fluorescence intensity values are plotted against GFP fluorescence intensity values for each animal across representative experimental conditions (n=380). Scale bar, 5 μm. **c**, Wild-type animals bearing integrated *ASI::FLP-7mCherry* and *CLM::GFP* transgenes were subjected to fasting at the indicated O_2_ levels, chosen because of URX activity state (ON at 21%; OFF at 10%)^22, 74^. The intensity of FLP-7mCherry fluorescence within a single coelomocyte was quantified and normalized to the area of CLM::GFP expression for each group. Data are expressed as a percentage of the normalized FLP-7mCherry fluorescence intensity of wild-type fed animals ± SEM (n=16-19). **p<0.01, ***p<0.001 by one-way ANOVA (Sidak). **d**, Wild-type animals bearing integrated *ASI::FLP-7mCherry* and *CLM::GFP* transgenes, along with indicated *ADL::HisCl* transgene. The insect HisCl channel is stimulated by histamine and inhibits neurons in which it is expressed^75^. The intensity of FLP-7mCherry fluorescence within a single coelomocyte was quantified and normalized to the area of CLM::GFP expression for each group. Data are expressed as a percentage of the normalized FLP-7mCherry fluorescence intensity of vehicle-treated non-transgenic animals ± SEM (n=26-48). ***p<0.001 by two-way ANOVA (Sidak). **e**, For intestine-specific RNAi, *sid-1; Pvha-6::sid-1* animals bearing integrated *ASI::FLP-7mCherry* and *CLM::GFP* transgenes were treated with vector, *pod-2, sbp-1 or elo-2* RNAi, the intensity of FLP-7mCherry fluorescence within a single coelomocyte was quantified and normalized to the area of CLM::GFP expression. Data are expressed as a percentage of the normalized FLP-7mCherry fluorescence intensity of animals treated with vector RNAi ± SEM (n=20-23). *p<0.05, **p<0.01, ***p<0.001 by one-way ANOVA (Dunnett).

**Extended Data Fig. 2.**
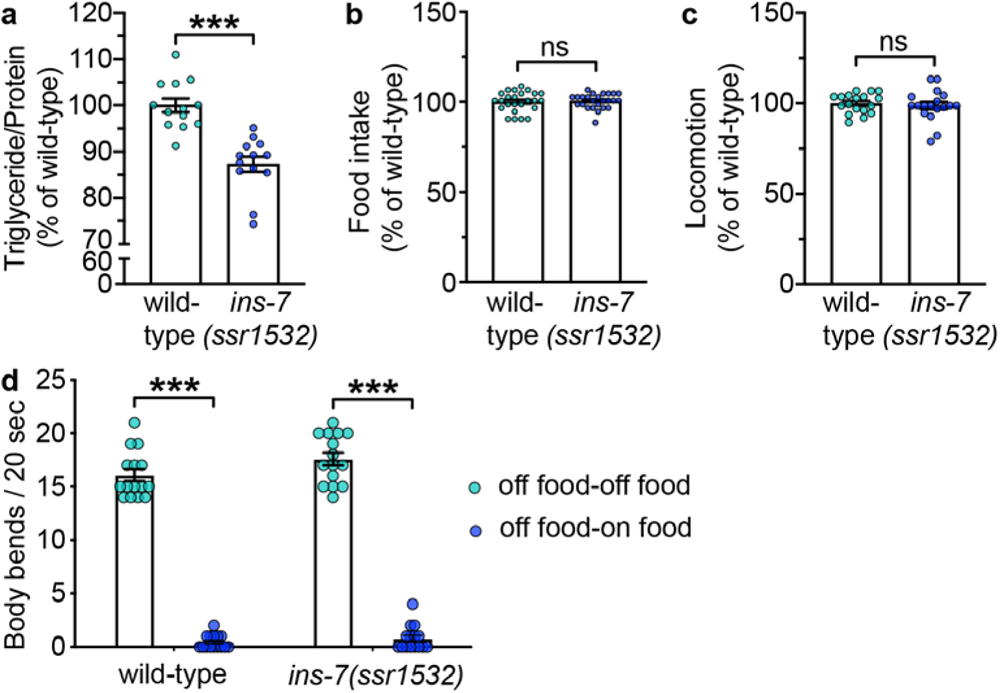
**a**, Triglyceride levels normalized to protein levels in wild-type and *ins-7(ssr1532)* mutants. Data are presented as a percentage of wild-type animals ± SEM (n=13). ***p<0.001 by Student’s *t*-test. Note that *ins-7(ssr1532)* mutants have decreased fat stores via Oil Red O measurements as well as triglyceride extraction, however the latter method extracts lipids from the whole animal including the germline and the embryos (which are lipid-rich), whereas the former method measures intestinal lipids only, a more faithful representation of the signaling axis under study. **b**, Food intake is expressed as a percentage of wild-type animals ± SEM (n=25). ^ns^p=0.7697 by Student’s *t*-test. **c**, Locomotion is expressed as a percentage of wild-type animals ± SEM (n=18). ^ns^p=0.5978 by Student’s *t*-test. **d**, For the enhanced slowing response assay, body bends per 20 seconds were measured in wild-type and *ins-7(ssr1532)* animals that were fasted (off food-off food) for 30min or animals that were fasted for 30min and then refed (off food-on food). Data are presented as Body bends/20 sec ± SEM (n=15). ***p<0.001 by two-way ANOVA (Sidak).

**Supplementary Table 1.**

| **Strains used in this study** | **Source** | **Identifier** | **Additional Information** |
| --- | --- | --- | --- |
| *N2*; *ssrls496*[*Patgl-1::GFP*] | PMID: 24120942 | SSR647 | Integrated. Backcrossed 7X. |
| *flp-7(ok2625)* | PMID: 28128367 | SSR1027 | Backcrossed 6X. |
| *N2*; *ssrls919*[*Pdaf-7::flp-7mCherry*]; *ssrls615*[*Punc-122::GFP*] | PMID: 28128367 | SSR1164 | Integrated. Backcrossed 6X. |
| *aak-2(ok524)*; *ssrls919*[*Pdaf-7::flp-7mCherry*]; *ssrls615*[*Punc-122::GFP*] | PMID: 28128367 | SSR1327 |  |
| *ins-7(tm1907)*; *ssrls919*[*Pdaf-7::flp-7::mCherry*]; *ssrls615*[*Punc-122::GFP*] | This paper | SSR1444 |  |
| *ins-7(tm1907)* | NBRP(Japan) | SSR1524 | Backcrossed 6X. |
| *sid-1(qt9)*; *ssrls919*[*Pdaf-7::flp-7mCherry*]; *ssrls615*[*Punc-122::GFP*]; *ssrls1218*[*Pvha-6::sid-1::GFP*] | This paper | SSR1541 | Backcrossed 4X. |
| *ins-7(ssr1532)*; *ssrls919*[*Pdaf-7::flp-7mCherry*]; *ssrls615*[*Punc-122::GFP*]; *ssrEx1192*[*Pges-1deltaB::ins-7::GFP*] | This paper | SSR1559 |  |
| *daf-16(mu86)*; *ssrls919*[*Pdaf-7::flp-7mCherry*]; *ssrls615*[*Punc-122::GFP*] | This paper | SSR1587 |  |
| *daf-2(e1370)*; *ssrls919*[*Pdaf-7::flp-7mCherry*]; *ssrls615*[*Punc-122::GFP*] | This paper | SSR1596 |  |
| *daf-2(e1370)*; *daf-16(mu86)*; *ssrls919*[*Pdaf-7::flp-7mCherry*]; *ssrls615*[*Punc-122::GFP*] | This paper | SSR1597 |  |
| *daf-2(e1370)*; *ueEx444*[*str-3::daf-2 sense:sl2mcherry, elt-2::gfp*]; *ssrls919*[*Pdaf-7::flp-7mCherry*]; *ssrls615*[*Punc-122::GFP*] | This paper | SSR1602 | From IV649: *daf-2(e1370)*; *ueEx444*[*str-3::daf-2 sense:sl2mcherry, elt-2::gfp*] |
| *ins-7(ssr1532)*; *ssrEx1143*[*Pins-7::ins-7::GFP*]; *ssrls919*[*Pdaf-7::flp-7mCherry*]; *ssrls615*[*Punc-122::GFP*] | This paper | SSR1607 |  |
| *ssrEx1272*[*Pstr-3::daf-2::GFP sense*]; *ssrEx1273*[*Pstr-3::daf-2::mcherry antisense*]; *ssrls919*[*Pdaf-7::flp-7mCherry*]; *ssrls615*[*Punc-122::GFP*] | This paper | SSR1616 |  |
| *N2*; *ssrls1240*[*Pges-1deltaB::ins-7mCherry*]; *ssrls615*[*Punc-122::GFP*] | This paper | SSR1617 | Backcrossed 6X. |
| *ins-7(ssr1532)*; *ssrls919*[*Pdaf-7::flp-7mCherry*]; *ssrls615*[*Punc-122::GFP*] | This paper | SSR1620 | Backcrossed 6X. |
| *ins-7(ssr1532)* | This paper | SSR1621 | Backcrossed 4X. |
| *ins-7(ssr1532)*; *daf-2(e1370)*; *ssrls919*[*Pdaf-7::flp-7mCherry*]; *ssrls615*[*Punc-122::GFP*] | This paper | SSR1622 |  |
| *aak-2(ok524)*; *daf-2(e1370)*; *ssrls919*[*Pdaf-7::flp-7mCherry*]; *ssrls615*[*Punc-122::GFP*] | This paper | SSR1625 |  |
| *ins-7(ssr1532)*; *ssrEx1192*[*Pges-1deltaB::ins-7::GFP*] | This paper | SSR1629 |  |
| *sid-1(qt9)*; *ssrls1220*[*Pges-1deltaB::sid-1::GFP*]; *ssrls1240*[*Pges-1deltaB::ins-7mCherry*]; *ssrls615*[*Punc-122::GFP*] | This paper | SSR1634 |  |
| *ins-7(ssr1532)*; *ssrEx1232*[*Pvha-6::ins-7::GFP*] | This paper | SSR1635 |  |
| *ins-7(ssr1532)*; *ssrEx1143*[*Pins-7::ins-7::GFP*] | This paper | SSR1640 |  |
| *ins-7(ssr1532); daf-16(mu86); Pdaf-16::daf-16GFP* | This paper | SSR1646 |  |
| *daf-16(mu86); Pdaf-16::daf-16GFP* | PMID: 18202391 | SSR1648 | From CF1934: *daf-16(mu86) l; muls109[daf-16p::gfp::daf-16 + odr-1p::rfp]*, Kenyon Lab |
| *ssrEx1295[Pstr-3::aak-2::GFP sense]; ssrEx1296[Pstr-3::aak-2::mcherry antisense]; ssrls919[Pdaf-7::flp-7mCherry]; ssrls615[Punc-122::GFP]* | This paper | SSR1651 |  |
| *ins-7(ssr1532); flp-7(ok2625)* | This paper | SSR1652 |  |
| *ins-7(ssr1532); ssrls496[Patgl-1::GFP]* | This paper | SSR1653 |  |
| *daf-2(e1370); daf-16(mu86); Pdaf-16::daf-16GFP* | This paper | SSR1661 |  |
| *N2; ssrls919[Pdaf-7::flp-7mCherry]; ssrls615[Punc-122::GFP]; ssrEx1192[Pges-1deltaB::ins-7::GFP]* | This paper | SSR1663 |  |
| *daf-2(e1370); ssrEx1295[Pstr-3::aak-2::GFP sense]; ssrEx1296[Pstr-3::aak-2::mcherry antisense]; ssrls919[Pdaf-7::flp-7mCherry]; ssrls615[Punc-122::GFP]; ssrEx1276[Plin-44::GFP]* | This paper | SSR1664 |  |
| *daf-16(mu86); Pdaf-16::daf-16GFP; ssrls1240[Pges-1deltaB::ins-7mCherry]; ssrls615[Punc-122::GFP]* | This paper | SSR1669 |  |

**Supplementary Table 2.**
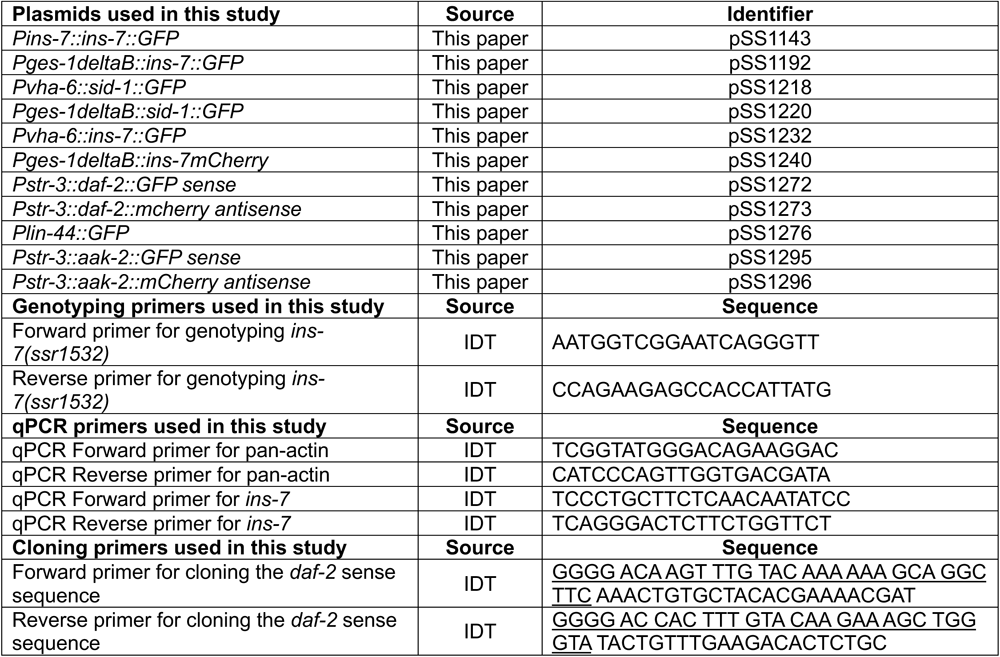
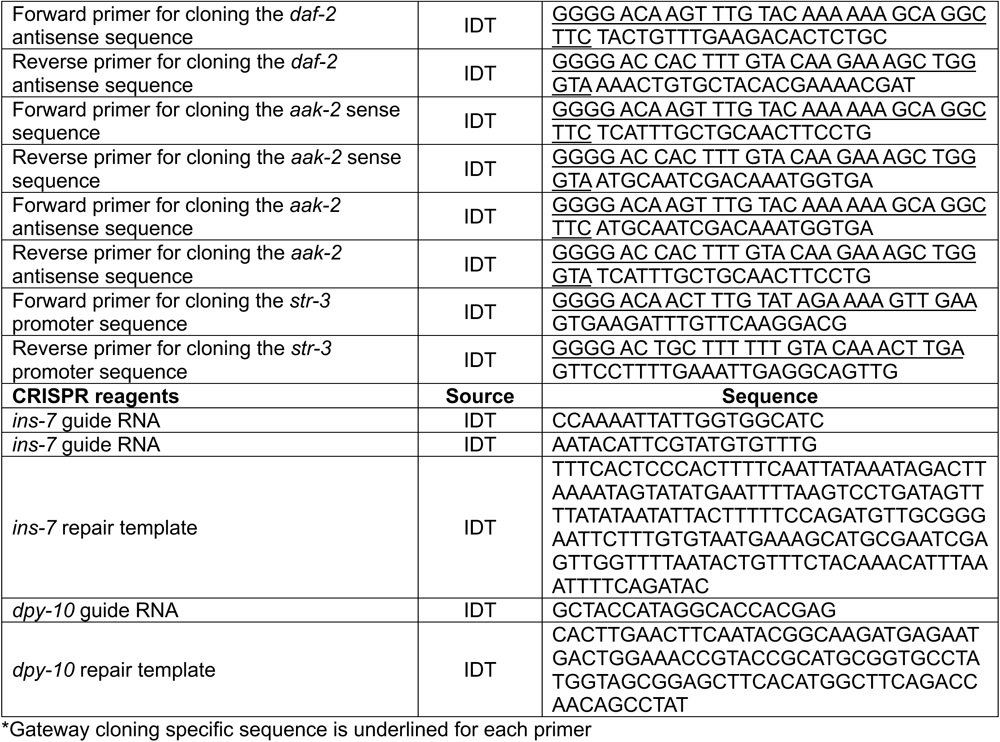

